# Differential roles of RIG-I-like receptors in SARS-CoV-2 infection

**DOI:** 10.1101/2021.02.10.430677

**Authors:** Duomeng Yang, Tingting Geng, Andrew G. Harrison, Penghua Wang

## Abstract

The retinoic acid-inducible gene I (RIG-I) and melanoma differentiation-associated protein 5 (MDA5) are the major viral RNA sensors that are essential for activation of antiviral immune responses. However, their roles in severe acute respiratory syndrome (SARS)-causing coronavirus (CoV) infection are largely unknown. Herein we investigate their functions in human epithelial cells, the primary and initial target of SARS-CoV-2, and the first line of host defense. A deficiency in MDA5 (*MDA5*^−/−^), RIG-I or mitochondrial antiviral signaling protein (MAVS) greatly enhanced viral replication. Expression of the type I/III interferons (IFN) was upregulated following infection in wild-type cells, while this upregulation was severely abolished in *MDA5*^−/−^ and *MAVS*^−/−^, but not in *RIG-I*^−/−^ cells. Of note, ACE2 expression was ~2.5 fold higher in *RIG-I*^−/−^ than WT cells. These data demonstrate a dominant role of MDA5 in activating the type I/III IFN response to SARS-CoV-2, and an IFN-independent anti-SARS-CoV-2 role of RIG-I.

## Introduction

Coronaviruses (CoV) are enveloped, positive sense single-stranded RNA viruses with the largest genomes (~30kb) among the known RNA viruses ^1^. They are the major etiologies of fatal human respiratory diseases, such as Severe Acute Respiratory Syndrome (SARS)-causing CoV and Middle East Respiratory Syndrome (MERS)-CoV. SARS started in November 2002 in Southern China, spread to 26 countries, and resulted in 8439 cases and 821 deaths^2^. Between its discovery in 2012 and January 2020, MERS-CoV had caused 2519 cases and 866 deaths^3^. At the end of 2019, a new SARS strain, SARS-CoV-2 that is 86% identical to SARS-CoV at the amino acid level, emerged in humans in Central China, and now has spread worldwide. Currently, there are >108 million confirmed SARS-CoV-2 cases and 2.4 million deaths from ~200 countries^4^, constituting the greatest global public health crisis in the 21^st^ century. Its pathogenesis, though plausibly similar to that of SARS-CoV, remains largely unknown – highlighting a critical need for new research in this area.

When invaded by a virus, a host cell produces a rapid innate immune response initiated by pathogen pattern recognition receptors (PRR), including C-type lectins, Toll-like receptors (TLR), retinoic acid-inducible gene I (RIG-I) like receptors (RLR), the cyclic GMP-AMP (cGAMP) synthase (cGAS), and nucleotide-binding oligomerization domain (NOD)-like receptors (NLR) etc. Once engaged by viral RNA, the cytoplasmic RLRs translocate and bind to a mitochondrion transmembrane protein, mitochondrial antiviral signaling protein (MAVS), which ignites a signaling event, leading to transcription of immune genes, in particular interferons (IFN) that provide an instant protection to the host ^5^. The relative contribution of different classes of PRRs to innate antiviral immune responses may vary with viral species and tissue cell types. The cytoplasmic RLRs (RIG-I and melanoma differentiation-associated protein 5, MDA5) are the essential PRRs to control RNA virus infection, while they show differential preference to different RNA viruses ^6^. MDA5 is essential for induction of the type I/III IFN response during mouse hepatitis virus (MHV), a murine coronavirus, infection in mice ^7,8^; while both RIG-I and MDA5 contribute to the type I IFN response in oligodendrocytes during MHV infection ^9^. Little is known about the differential role of RLRs in SARS-CoV infection, although they can be hijacked by SARS-CoV proteins to evade host immune responses ^10 11^.

## Results and discussion

We investigated the differential role of RIG-I and MDA5 in controlling SARS-CoV-2 infection and mounting immune responses in a human lung epithelial cell lines Calu-3. We generated individual RIG-I, MDA5 and MAVS knockout using CRISPR-Cas9 and validated by immunoblotting (**Fig.1a**). To prove that these gene function was precisely silenced, we infected mutant cells with vesicular stomatitis virus (VSV specifically activates RIG-I-MAVS) with an green fluorescence protein (GFP) integrated into its genome. As expected, *RIG-I*^−/−^ or *MAVS*^−/−^ cells presented a much higher VSV-GFP load than wild type (WT) cells, while *MDA5*^−/−^ cells had a similar viral load as WT (**Fig.1b**). These results demonstrate precise disruption of each gene function of interest by CRSIPR-Cas9. We then compared SARS-CoV-2 infection and interferon responses in these knockout cells in parallel. The intracellular viral RNA load was increased by ~2-3.5-fold in all knockout cells at 24hrs post infection (p.i.), by 5-12-fold at 72hrs p.i. (**Fig.2a**). Consistently the extracellular viral titers were also much higher from all knockout cells than WT (**Fig.2b**). We confirmed these observations in A549 cells (**Fig.2c**). Although primarily sensing DNA viruses, the cGAS-STING signaling pathway also restricts many RNA virus infection ^12,13^. We noted a slight increase in SARS-CoV-2 load in *STING*^−/−^ cells (**Fig.2 d, e**), suggesting that STING signaling is largely dispensable.

**Fig.1.**
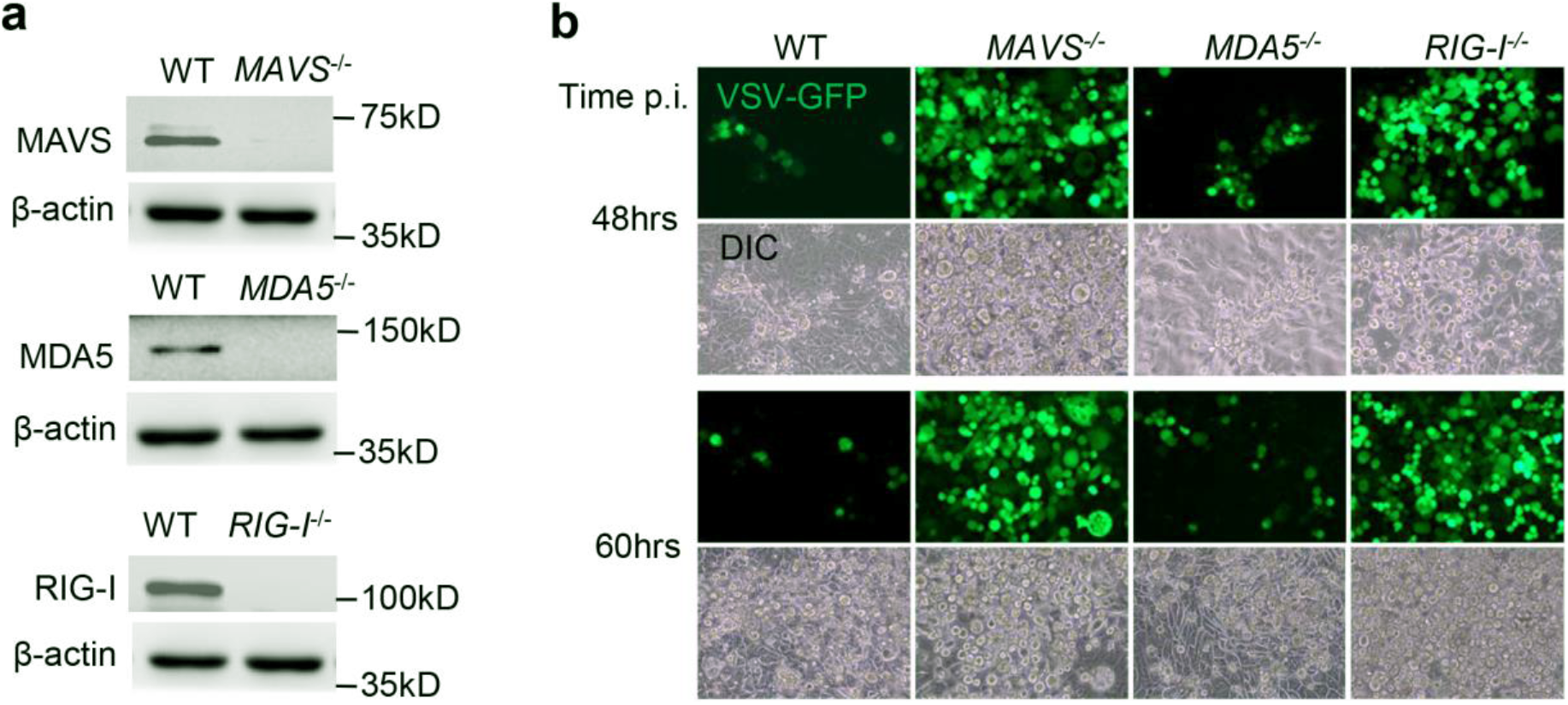
Functional validation of gene knockouts by CRISPR-Cas9. **a**) The immunoblots show gene knockout efficiency in Calu-3 cells. β-actin is a house keeping gene and serves as a protein loading control. **b**) Fluorescent microscopic images of VSV-GFP at several time points post infection (p.i.). Magnification: 100 x. DIC: differential interference contrast. The results are representative two reproducible independent experiments.

**Fig.2.**
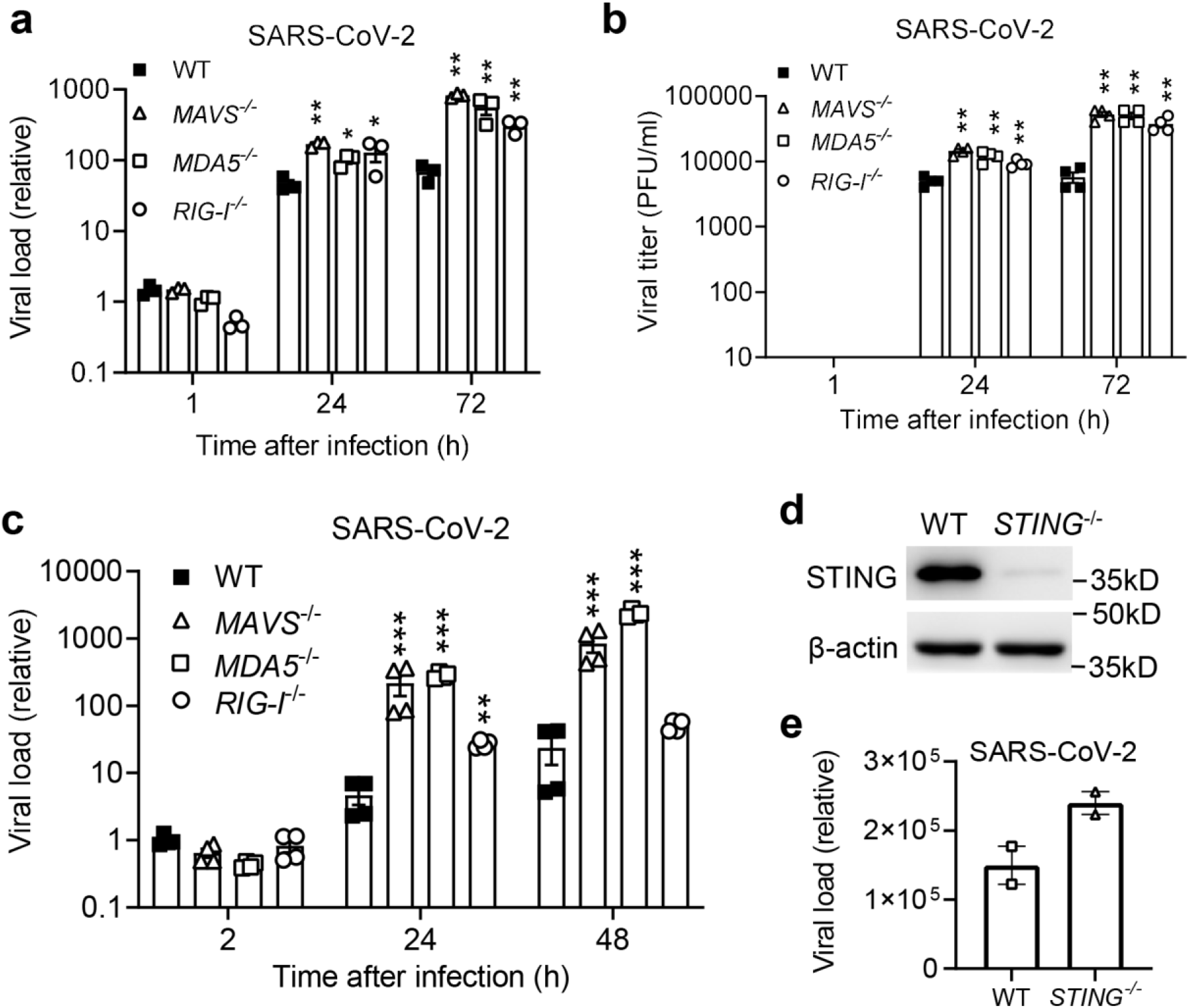
An essential role of the MDA5-MAVS axis in control of SARS-CoV-2 infection. **a**) Quantitative RT-PCR analyses of SARS-CoV-2 RNA loads in Calu-3 cells infected with SARS-CoV-2 at a multiplicity of infection (MOI) of 0.5. **b**) The extracellular viral titers in the cell culture supernatants of Calu-3 cells. PFU: plaque forming unit. **c**) Quantitative RT-PCR analyses of SARS-CoV-2 RNA loads in A549 cells infected with SARS-CoV-2 at a MOI of 0.5. **d**) The immunoblots show STING knockout efficiency in Calu-3 cells. β-actin is a house keeping gene and serves as a protein loading control. **e**) Quantitative RT-PCR analyses of SARS-CoV-2 RNA loads in Calu-3 cells infected with SARS-CoV-2 at a MOI of 0.5. Each symbol=one biological replicate. All the data are presented as mean ± S.E.M. and statistical significances are analyzed by unpaired two-tailed Student’s t-test, * *P*<0.05, ** *P*<0.01, ***, P<0.001. The results are representative two reproducible independent experiments.

We next examined antiviral immune responses. The *IFNB1* (type I IFN) and *IL29* (type III IFN) mRNA levels were continuously upregulated during the course of infection in WT cells; while this induction was impaired in *MDA5*^−/−^ and *MAVS*^−/−^cells, though not completely abolished; So was one of interferon-stimulated gene (ISG15) (**Fig.3a)**. The concentrations of IFN-β and C-X-C motif chemokine ligand 10 (CXCL10) proteins in the cell culture supernatants from *MDA5*^−/−^ and *MAVS*^−/−^ were much lower than WT cells (**Fig.3b**). However, the type I/III IFN and *ISG15* expression was higher in *RIG-I*^−/−^ than WT cells (**Fig.3a**), suggesting that RIG-I interferes with SARS-CoV-2 replication independently of IFNs. Of note, the angiotensin-converting enzyme 2 (ACE2, a major cell entry receptor for SARS-CoV-2) mRNA expression was continuously upregulated during the course of infection in WT cells, and it was ~2.5-fold higher in *RIG-I*^−/−^ than that in WT other knockout cells (**Fig.3c**), suggesting that RIG-I represses ACE2 mRNA expression.

**Fig.3.**
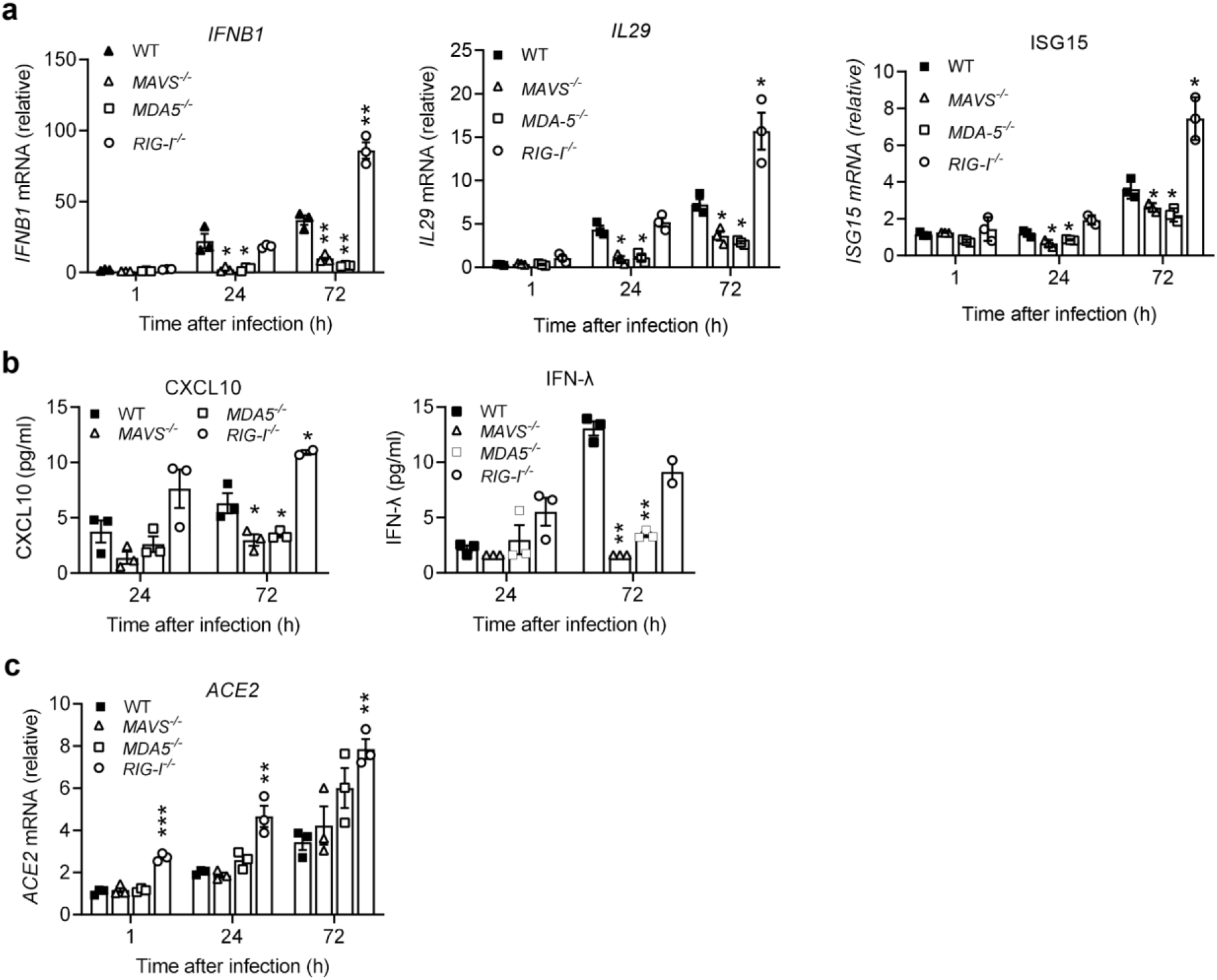
An essential role of the MDA5-MAVS axis in induction of type I/III IFNs by SARS-CoV-2. **a**) Quantitative RT-PCR analyses of immune gene transcripts and **b**) Quantification of IFN-λ and CXCL10 proteins by ELISA, in Calu-3 cells infected with SARS-CoV-2 at a multiplicity of infection (MOI) of 0.5. **c**) Quantitative RT-PCR analyses of ACE2 mRNA. All the data are presented as mean ± S.E.M. and statistical significances are analyzed by unpaired two-tailed Student’s t-test, * *P*<0.05, ** *P*<0.01, ***, P<0.001. Each symbol=one biological replicate. The results are representative two reproducible independent experiments.

Understanding the major PRR pathways in the respiratory tract epithelial cells is physiologically meaningful as these cells are the primary and initial target of SARS-CoV-2, and the first line of host defense. The RLR signaling is functional in all tissues and cell types, in contrast to viral RNA-sensing TLRs (TLR3, TLR7) that are primarily limited to immune cells. Thus, herein we only focused on RLRs in lung epithelial cells. Our results demonstrate that MDA5 is the predominant RLR, because MDA5 deficiency led to a similar effect on viral replication and type I/III IFNs as MAVS deletion did (**Fig.2, 3**). However, in neither MDA5 nor MAVS knockout cells, induction of IFNs was completely abolished (**Fig.3**), suggesting that other PRR signaling pathways may collectively play a role. Surprisingly, RIG-I deletion had no negative impact on induction of IFN responses by SARS-CoV-2, but still increased viral replication (**Fig.2, 3**), suggesting that RIG-I plays a MAVS-IFN-independent antiviral role. Of note, ACE2 expression was upregulated by ~2.5 folds in *RIG-I*^−/−^, compared to WT and other knockout cell even at 1hr post infection. This suggests RIG-I may restrict SARS-CoV-2 cellular entry by repressing ACE2 mRNA expression. Alternatively, RIG‐I may exert broad antiviral activity in the models that have defective IFN signaling^14^. RIG-I could induce expression of antiviral effectors by an IFN-Janus kinase (JAK)-signal transducer and activator of transcription (STAT)-independent mechanism ^15^. RIG-I could disrupt interaction of hepatitis B virus polymerase with the 5’-ε region of viral pregenomic RNA in an RNA-binding dependent manner, which consistently suppressed viral replication ^16^. RIG-I binding to the nucleocapsid of influenza A virus could directly inhibit viral replication ^17^. Both MDA5 and RIG-I may displace viral proteins pre-bound to dsRNA in a manner dependent on their ATP hydrolysis, thus exerting a MAVS-IFN-independent broad-spectrum antiviral role (Sindbis, yellow fever, Coxsackie B, Sendai, influenza A, and human parainfluenza 3 viruses) ^18^. During the submission of this manuscript, a study showed an important anti-SARS-CoV-2 role of MDA5 that is consistent with our results, however dispensable function of RIG-I in limiting SARS-CoV-2 infection^19^. This discrepancy could be due to a difference in the cell line and gene knockout method.

In summary, we report a dominant role of MDA5 in sensing SARS-CoV-2 and activating immune responses, and an IFN-independent anti-SARS-CoV-2 role of RIG-I. Given a therapeutic potential of IFN in COVID-19 ^20 21^ and the essential role of MDA5 in IFN induction, the MDA5 agonists could thus be therapeutic against SARS-CoV-2 infection.

## Materials and Methods

### Antibodies, cells and viruses

The rabbit anti-MDA5 (Cat# 5321), RIG-I (Cat# 3743), and Actin (Cat# 8456) were purchased from Cell Signaling Technology (Danvers, MA, United States). Mouse anti-human MAVS (Cat# SC-365333) was a product of Santa Cruz Biotechnology (Santa Cruz, CA, United States). Human embryonic kidney (HEK) 293 T (Cat# CRL-3216), Vero cells (monkey kidney epithelial cells, Cat# CCL-81), human lung epithelial A549 (Cat# CCL-185), human lung epithelial Calu-3 (Cat# HTB-55) cell lines were from American Type Tissue Culture (Manassas, VA, United States). SARS-CoV-2 (NR-52281, Isolate USA-WA1/2020) was provided by BEI Resources (funded by National Institute of Allergy and Infectious Diseases and managed by ATCC, United States). The vesicular stomatitis virus (VSV) used in this study was Indiana strain, and the green fluorescence protein (GFP) tagged VSV was provided by Dr. Rose at Yale University^22^. These viruses were propagated in Vero cells.

### Cell culture and virus infection

HEK293T/Vero cells and Calu-3/A549 cells were grown in Dulbecco’s modified Eagle’s medium or Roswell Park Memorial Institute (RPMI) 1640, respectively (Life Technologies, Grand Island, NY, USA) supplemented with 10% fetal bovine serum (FBS) and antibiotics/antimycotics. These cell lines are not listed in the database of commonly misidentified cell lines maintained by ICLAC, and have not been authenticated in our hands. They were routinely treated with MycoZAP (Lonza) to prevent mycoplasma contamination.

SARS-CoV-2/VSV-GFP was added to cells at a multiplicity of infectivity (MOI) of 0.5 respectively, and incubated at 37°C, 5% CO_2_ for 1-2hrs. The viral inoculum was then removed completely, and replaced by fresh culture medium. The cells and medium were collected immediately (1hr or 2hrs as a baseline), or 24-72hrs after inoculation.

### Gene knockout by CRISPR-Cas9

A gene specific guide RNA was cloned into lentiCRISPR-V2 vector and co-transfected into HEK293T cells with the packaging plasmids pVSV-G and psPAX2. Forty-eight hours after transfection, the lentiviral particles in the cell culture media were applied to A549 or Calu-3 cells for 48 hours. The transduced cells were then selected with puromycin at 2μg/ml for 4-5 days until non-transfected control cells completely died. The guide RNA for human RIG-I, MDA5 and MAVS was TCCTGAGCTACATGGCCCCC, CTTTCTGCCTGCAGAGGTGA, and AAGTTACCCCATGCCTGTCC respectively ^23,24^. The wild type control was lentiCRISPRv2 vector only.

### Plaque forming assay

Quantification of infectious viral particles in cell culture supernatant was performed on Vero cell monolayer ^25^. Briefly, serial dilutions of supernatants was applied to confluent Vero cells (6-well plate) at 37°C for 2 h. The inoculum was then removed and replaced with 2 ml of DMEM complete medium with 1% SeaPlaque agarose (Cat# 50100, Lonza). Plaques were visualized using Neutral red (Sigma-Aldrich) after 3 days of incubation at 37°C, 5% CO_2_.

### Reverse transcription and quantitative (q)PCR

Up to 1×10^6^ culture cells were collected in 350 μl of RLT buffer (QIAGEN RNeasy Mini Kit). RNA was extracted following the QIAGEN RNeasy manufacturer’s instructions. Reverse transcription of RNA into complementary DNA (cDNA) was performed using the BIO-RAD iScript™ cDNA Synthesis Kit. Quantitative PCR (qPCR) was performed with gene-specific primers and SYBR Green PCR master mix. Results were calculated using the –ΔΔCt method and a housekeeping gene, beta actin, as an internal control. The qPCR primers and probes for immune genes were reported in our previous studies^24^. The primers for SARS-CoV-2 were forward: 5’-GAC CCC AAA ATC AGC GAA AT-3’ and reverse: 5’-TCT GGT TAC TGC CAG TTG AAT CTG-3’.

### Multiplex enzyme-linked immunosorbent assay (ELISA)

We used a LEGENDPlex (Biolegend, San Diego, CA 92121, USA) bead-based immunoassay to quantify the cytokine concentrations in the cell culture supernatant of SARS-CoV-2 infected cells. The procedures were exactly the same as in the product manual. Briefly, the supernatants or standards were mixed with antibody-coated microbeads in a filter-bottom microplate, and incubated at room temperature for 2hrs with vigorous shaking at 500 rpm. After removal of unbound analytes and two washes, 25 μL of detection antibodies were added to each well, and the plate was incubated at room temperature for 1hr with vigorous shaking at 500 rpm. 25 μL of SA-PE reagent was then added directly to each well, and the plate was incubated at room temperature for 30min with vigorous shaking at 500 rpm. The beads were washed twice with wash buffer, and then transferred to a microfuge tube. The beads were fixed with 4% PFA for 15min and resuspended in FACS buffer. The beads were run through a BIORAD ZE5 and the concentrations of analytes were calculated with the standards included using the LEGENDPlex software.

## Acknowledgements

This work was supported in part by a National Institutes of Health grant R01AI132526 to P.W.

## Author contributions

D.Y. performed the majority of the experimental procedures and data analyses. T.G. and A.G.H. contributed to some of the figures. P.W. conceived and oversaw the study. D.Y. and P.W. wrote the paper and all the authors reviewed and/or modified the manuscript.

## Conflict of Interest

The authors declare no competing financial/non-financial interest.

## Notes

### Competing Interest Statement

The authors have declared no competing interest.

